# High Resolution Repli-Seq defines the temporal choreography of initiation, elongation and termination of replication in mammalian cells

**DOI:** 10.1101/755629

**Authors:** Peiyao A. Zhao, Takayo Sasaki, David M. Gilbert

## Abstract

DNA replication in mammalian cells occurs in a defined temporal order during S phase, known as the replication timing (RT) programme. RT is developmentally regulated and correlated with chromatin conformation and local transcriptional potential. Here we present RT profiles of unprecedented temporal resolution in two human embryonic stem cell lines, human colon carcinoma line HCT116 as well as F1 subspecies hybrid mouse embryonic stem cells and their neural progenitor derivatives. Strong enrichment of nascent DNA in fine temporal windows reveals a remarkable degree of cell to cell conservation in replication timing and patterns of replication genome-wide. We identify 5 patterns of replication in all cell types, consistent with varying degrees of initiation efficiency. Zones of replication initiation were found throughout S phase and resolved to ~50kb precision. Temporal transition regions were resolved into segments of uni-directional replication punctuated with small zones of inefficient initiation. Small and large valleys of convergent replication were consistent with either termination or broadly distributed initiation, respectively. RT correlated with chromatin compartment across all cell types but correlations of initiation time to chromatin domain boundaries and histone marks were cell type specific. Haplotype phasing revealed previously unappreciated regions of allele-specific and allele-independent asynchronous replication. Allele-independent asynchrony was associated with large transcribed genes that resemble common fragile sites. Altogether, these data reveal a remarkably deterministic temporal choreography of DNA replication in mammalian cells.

- Highly homogeneous replication landscape between cells in a population
- Initiation zones resolved within constant timing and timing transition regions
- Active histone marks enriched within early initiation zones while enrichment of repressive marks is cell type specific.
- Transcribed long genes replicate asynchronously.

## INTRODUCTION

DNA replication in eukaryotes proceeds in a defined temporal order known as the replication timing (RT) programme (Rhind and Gilbert, 2013). RT is dynamically regulated during development and differentiation and exhibits cell-type specific RT signatures (Hiratani et al., 2008; Rivera-Mulia et al., 2015). RT is also closely correlated with the A/B compartment of chromatin structure, local chromatin environment and the transcription potential of the region (Moindrot et al., 2012; Pope et al., 2014; Sima et al., 2019). One of the primary assays for genome-wide RT has been E/L Repli-Seq in which cells labelled with BrdU for 10-20% of S phase are sorted into early and late S fractions and RT profiles are generated from the log2 ratio of read enrichment in the BrdU-immunoprecipitated early fraction to that in the late fraction (E/L) (Marchal et al., 2018). E/L RT profiles reveal large Constant Timing Regions (CTRs) of early and late replication manifesting as plateaus and punctuated by Timing Transition Regions (TTRs) of rightward or leftward slopes (Dileep et al., 2015). Early and late CTRs must contain sites of replication initiation as they replicate too rapidly to be accounted for by elongation alone. By contrast, TTRs are hypothesised to consist mainly of unidirectional forks, occasionally accelerated by sequentially firing origins (Farkash-Amar et al., 2008; Guilbaud et al., 2011; Hiratani et al., 2008). However, the measurements supporting these hypotheses are derived from E/L Repli-Seq, which smooths data over hundreds of kilobases (kbs) and so lacks the resolution to identify sites of replication initiation or predict the presence or absence of origins in TTRs. Moreover, since the E/L method averages all stochastic variation in a cell population, it does not permit one to determine how variable the RT of any given chromosome segment might be. We have developed a single cell Repli-Seq method to address this problem, concluding that the RT program is stable from cell to cell, but this method still suffers from low resolution (Dileep and Gilbert, 2018; Takahashi et al., 2019).

Mammalian replication origin mapping poses a significant challenge due to the high flexibility of sites that can initiate replication and their varying efficiencies (Gilbert, 2001; Hyrien, 2015). Moreover, frequently clusters of origins used at varying efficiencies produce what have been called “initiation zones”. Small nascent strand sequencing (SNS-seq) and Okazaki fragment sequencing (OK-seq) methods have produced comprehensive maps of replication origins and fork polarities, respectively, in several human and mouse cell lines (Besnard et al., 2012; Cayrou et al., 2015; Fu et al., 2014; Petryk et al., 2018; Petryk et al., 2016). SNS can map sites to kilobase resolution but the sites detected must fire frequently enough to detect above noise; clusters of inefficient origins escape detection. OK-seq detects transitions in fork polarity that define bi-directional replication but at low (>50kb) resolution, detecting the sum efficiency of initiation across initiation zones. Mapping origins on single DNA molecules can, in principle, measure the frequency of initiation at specific sites but existing methods are extremely low throughput. Studies analyzing hundreds of DNA fibres from a single genomic location have revealed that some regions initiate at defined sites (Anglana et al., 2003; Demczuk et al., 2012) while other genomic locations can initiate at many sites distributed broadly, with each site used in less than 2% of S phases (Anglana et al., 2003; Demczuk et al., 2012). By contrast, RT shows little cell to cell variation (Dileep and Gilbert, 2018; Takahashi et al., 2019) Thus, the prevailing view is that a deterministic RT programme emerges from stochastic origin selection (Rhind et al., 2010). However, this hypothesis still relies on low throughput, low resolution methods; higher-throughput and more sensitive methods are needed to understand the landscape of origin and fork distributions genome wide in mammalian cell.

To address these gaps, we developed a high resolution Repli-Seq approach that can delineate initiation zones (IZs), TTRs and termination sites with unprecedented temporal resolution in 3 human cell lines and 2 mouse cell types. We identify specific sites of previously undetected replication initiation activity within TTRs and resolve them from stretches of uni-directional replication or true TTRs. We detect a remarkable homogeneity in the temporal order of replication in cell populations, at considerably higher resolution than achieved by our prior single cell measurements (Dileep and Gilbert, 2018). Whereas active histone modifications were consistently correlated with early firing IZs, repressive histone marks varied between cell types in their relationship with IZ initiation time. However, initiation time was intimately linked to Hi-C compartment and IZs were enriched at Hi-C domain boundaries. The temporal resolution permitted the identification of bi-phasically replicated regions and the extensive read depth and breadth permitted haplotype phasing, which revealed both allele-specific replication asynchrony and allele-independent asynchrony, the latter of which was associated with long transcribed genes that have been associated with common fragile sites (CFS).

## RESULTS

### 16 fraction Repli-seq reveals patterns of replication with high temporal resolution

We performed high resolution Repli-Seq in 5 cell types: three human cell lines, male and female human embryonic stem cell (hESC) lines H1 (WA01) and H9 (WA09) hESCs and human colorectal cancer line HCT116 as well as mouse embryonic stem cell (mESC) line F121-9 derived from hybrid *castaneusXmusculus* mouse embryos and finally neural progenitor cells (mNPCs) derived from F121-9. Cells were labelled with BrdU for 30 mins, stained with propidium iodide and sorted by FACS into 16 equal S phase fractions (**Fig1a**). 16 fractions were chosen due to its approximate equivalence to the fraction of S phase labelled with BrdU (30min of an 8-10 hour S phase) as well as convenience with 4-way FACS sorting. BrdU-immunoprecipitated DNA from each fraction was validated by qPCR on regions whose RT is known from prior E/L Repli-Seq data (**supplementary Fig1**). The raw read counts per 50kb bin for each S phase fraction were corrected for mappability using G1 whole genome sequencing, each 50 kb bin was then Gaussian smoothed (**Methods, supplementary figure 2**) and scaled. Repli-Seq heatmaps for all cell lines assayed were visually inspected to ensure that features such as diffuse peaks in the heatmap, which represent less efficient IZs, were preserved. Datasets were subsequently normalised between fractions by scaling column-wise between bins so that the sum for any individual column amounted to 100 (Methods, **Fig1a**). High-Resolution Repli-Seq heatmaps show high concordance with E/L Repli-Seq while providing insights into the finer features of replication that we will further expound in the next section (**Fig1b-d**). Consistent with the prior knowledge that RT profiles are cell type specific, we find high genome-wide similarity between H1 and H9 16 fraction datasets while the HCT116 dataset shows less correlation with hESCs (**Supplementary Fig3**). Mouse heatmaps are generated for both maternal *musculus (mus)* and paternal *castaneous (cas)* genomes after allele parsing. The two genomes in mESCs and those in mNPCs show high correlation within the same cell type and lower correlation across cell types. (**Supplementary Fig4**).

**Figure 1.**
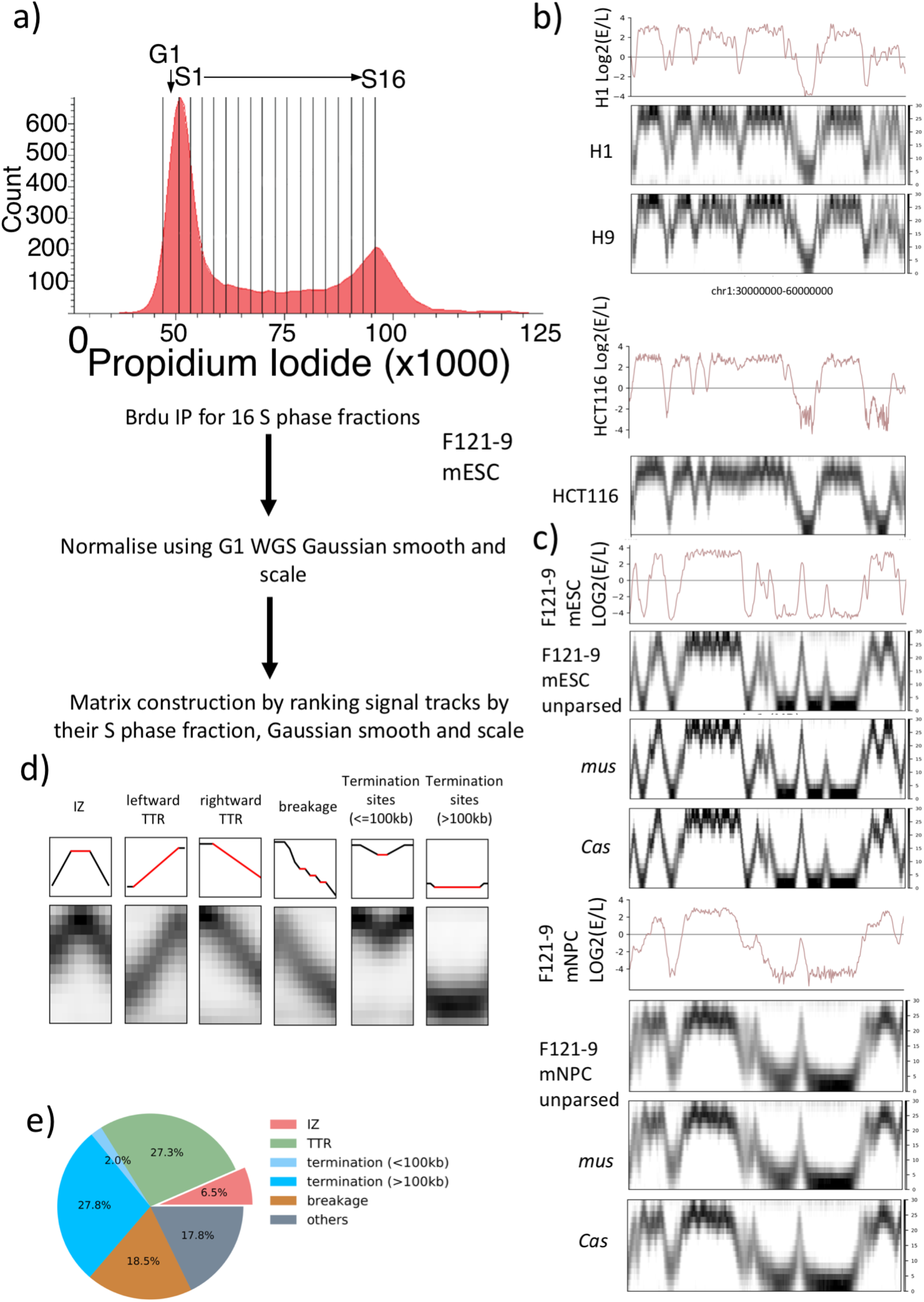
High-Resolution Repli-Seq produces robust and reproducible heatmaps that annotate features of replication at fine temporal resolution. a) Experimental and analysis flow for High-Resolution Repli-seq. b) E/L Repli-seq for H1 hESCs (top), normalised High-Res Repli-seq heatmaps for chr1:30000000-60000000 in H1, H9, HCT116. c) E/L Repli-seq for F121-9 mESCs and mNPCs, I normalised High-Res Repli-seq heatmaps for chr1:125000000-155000000 in unparsed *mus* allele, *cas* allele in mESC and mNPC. d) Features observed in High-Resolution Repli-Seq heatmap. Example schematics (top row: red indicates the corresponding feature and black indicates the surrounding bins) and corresponding heatmaps (bottom row) of IZs, leftward TTR, rightward TTR, breakages in TTR, short termination sites (<=100kb) and long termination sites (>=100kb). e) Percentage of genome constituted by features in d).

### Defining features in Repli-seq heatmaps: IZs, TTRs, TTR breakages and termination sites

Using high-resolution Repli-Seq we were able to identify 5 distinct features in the landscape of DNA replication across the genome (**Fig1d**). Since origins are replicated earlier than their surrounding regions, and since the window size is 50kb and cannot distinguish single highly efficient origins from clusters of less efficient origins, we defined “initiation zones” (IZs) as sites manifesting as vertical peaks in the heat maps. Emanating from IZs are segments of constant slope traversing through several S phase temporal windows, which represent the TTRs, presumably consisting of replication forks moving away from initiation sites during the progression of S phase. Depending on the directionality of progression of the forks emerging from IZs, in **Fig. 1d** TTRs are illustrated as either leftward or rightward TTRs (ascending or descending) but are considered a single feature of the data. Frequently found within TTRs are small decreases in slope (hereafter referred to as ‘breakages’) that (see below) signify origin firing accelerating the rate of DNA replication at these regions. V-shaped features represent defined sites of replication termination (<100kb) where opposing replication forks fuse. Finally, large U-shaped termination sites (>100kb) are too large and synchronously replicated to be explained by fork fusion and therefore must contain replication origins (Conti et al., 2007). While IZs constitute less than 10% of the genome (**Fig 1e**), more than half of the genome consists of breakages or large TTRs, indicating that 72% of the genome has detectable replication initiation potential. Origin-free regions, which constitute TTRs and termination sites (<100kb), constitute ~28% of the genome. Altogether, these results provide a comprehensive view of the kinetics of DNA replication genome wide conserved in several mammalian cell types.

We identified ~3000 IZs in human cell lines and ~2200 IZ in mouse cell lines. We classified IZs into early (S1-3), early-mid (S4-6), late-mid (S7-9) and late S (S10-12) IZs depending on the S phase fractions where the highest read density was identified (see methods). Pile up heatmap images centred on each of these temporally defined IZs in each cell type are shown in **Fig2a**. In both human and mouse, we identified constitutive IZs that are shared between cell types as well as cell-type specific IZs (**Fig2b**). A significant proportion of IZs are shared between cell types (1933 in human cell lines and 991 in mouse for IZs shared between cell types as well as between alleles). H1 and H9 share an intersecting set of 670 IZs that are unique to hESCs while 746 IZs are unique to HCT116. mESC *mus* and *cas* alleles share 650 mESC-specific IZs while mNPC *mus* and *cas* alleles share 464 mNPC-specific IZs.

**Figure 2.**
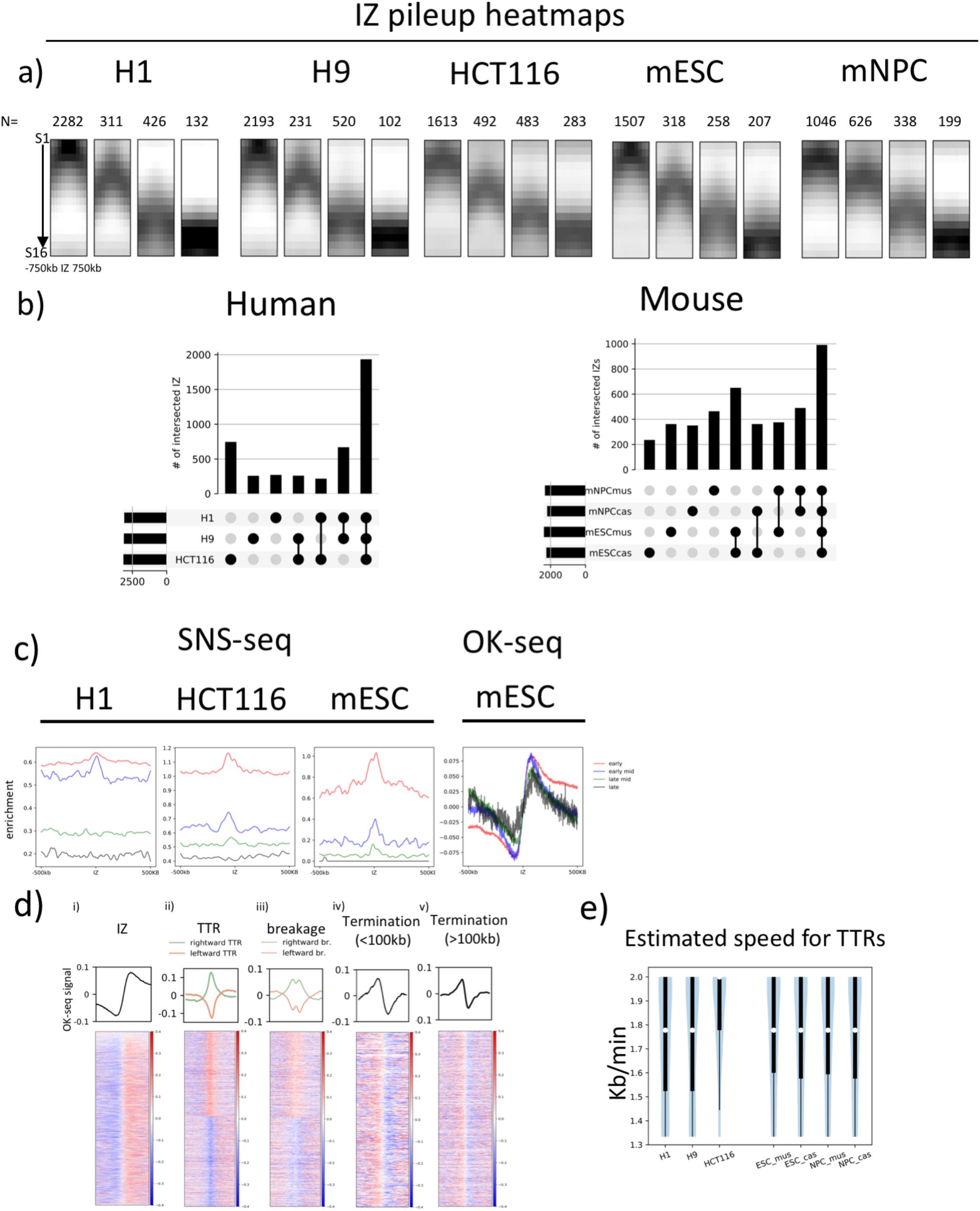
High Resolution Repli-Seq identifies features highly concordant with Ok-seq. a) Pile ups of IZs that are categorised into four categories: early, early-mid, I late-mid and late by the timing of initiation in H1 and H9 hESCs, HCT116, F121-9 mESCs and mNPCs. The number of IZs in each category is shown on top of the pile up heatmap. b) UpSet plots showing numbers of IZs either unique to each cell type alone or uniquely in common between the connected cell types in human and mouse cell lines. Black bars on the lower left are total IZs in each cell type. c) Mean line plots showing SNS-seq signal and OK-seq signal centred at IZs of early (red), early-mid (blue), late-mid (green) and late (black) RT +/− 500kb. d) Mean line plots (top row) and heatmaps (bottom row) showing OK-seq signal centred at IZs (i), TTRs (ii), breakages (iii), termination sites (<100kb) (iv) and termination sites(>100kb) (v) where negative values indicate an enrichment for Okazaki fragments at leftward fork and positive values indicate an enrichment for Okazaki fragments at rightward fork. Each row in a heatmap represent a single site of any feature. Heatmaps are sorted by row using site size. TTR (ii) and breakage (iii) heatmaps are arranged so that rightward TTR/breakages are stacked on top of leftward TTR/breakage and sorted separately. Green and orange mean line plots represent the rightward and left lef-tward TTRs respectively. e) Distribution of estimated TTR speed in kb/min for H1, H9, HCT116, mESC and mNPC. White dots represent the medians of distribution. Thick black lines mark the distribution between the first quartile to the third quartile. Thin black lines indicate the complete distribution of TTR speeds.

Next, we aligned our IZs with IZs and initiation sites identified through other origin mapping methods applied in these cell lines, namely small nascent strand sequencing (SNS-seq) and Okazaki fragment sequencing (OK-seq). For H1, HCT116 and the *musculus* genome in mESCs, the SNS-seq shows an enrichment of SNS signal centred around early IZs but not late IZs (**Fig2c**). Moreover, the magnitude of SNS-seq signal within and surrounding the IZs decreases as the timing of initiation within IZs becomes later in S phase. The trend was true across all human and mouse cell types examined, although quantitative cell type-specific differences exist. On the other hand, IZs show the expected transition in fork polarity that signifies the presence of initiation sites in Ok-seq dataset of the same cell type (mESCs) for all IZs regardless of their S phase initiation time (**Fig2c**). Even the very late IZs that are devoid of SNS-seq origin signal correspond to regions of OK-seq polarity transition showing that OK-seq and High-Resolution Repli-Seq validate each other and identify IZs of the same dynamic characteristics. Reciprocally by centring on all OK-seq IZs called in (Petryk et al., 2018) and plotting the pile-up of High-Resolution Repli-Seq signal, we show that called OK-seq IZs are primarily early IZs (**Supplementary Fig5**). Thus, the earliest initiating IZs are the most efficient at generating bidirectional replication forks, but all IZs do so. The discrepancy between OK-seq and SNS-seq is likely due to the fact that the two methods enrich for initiation events of differing properties. SNS-seq primarily identifies the most efficient origins that fire within approximately 1 kb and these have been shown to fire in early S phase (Gilbert, 2010; Hyrien, 2015), hence the prevalence of SNS-seq signal around early IZs. OK-seq is lower resolution and hence does not detect most isolated origins due to the fact that even the high efficiency SNS origins are still firing in a percentage of cells too low to skew Okazaki fragment distribution. However, OK-seq is good at detecting clusters of origins that sum up to a high efficiency zone of initiation and therefore manifest as Okazaki fragment strand switches within these zones. This is true whether the origins are high efficiency or low efficiency so long as the sum total efficiency is sufficient to create a detectable OK-seq strand switch, thus such switches can be found at all times during S phase. Overall, we conclude that High-Resolution Repli-Seq provides an orthogonal confirmation of the initiation zones identified by OK-seq mapping with temporal resolution. The presence of SNS-seq signal within the earliest replicating IZs is consistent with the presence of efficient origins within this class of IZs.

Due to the high consistency between Repli-Seq IZs and OK-seq signal, we examined OK-seq signal centred around all of the dynamic features of replication described in **Fig1d**. TTRs were divided into either rightward or leftward TTRs and associated with positive or negative OK-seq signal values respectively, consistent with unidirectional fork movement (**Fig2dii)**. In addition, SNS-seq signal around rightward or leftward TTRs reveals a sharp drop in signal enrichment from the IZ end to the termination site end of the TTR depending on its orientation, indicating that TTRs are indeed depleted of SNS-detected origins and segregate IZs that are rich in efficient origins from termination sites that contain either no origins or inefficient origins (**Supplementary Fig6**). TTR breakages and large termination sites (>100kb) are expected to contain origin activity to replicate so rapidly (albeit small ones could, in principle, be regions of unusually rapid fork movement). In fact, TTR breakages were detected as a drop in fork polarity in either rightward or leftward fork movement depending on the directionality of the TTR in which the breakage is located, consistent with origin activity causing diminished polarity bias of Okazaki fragments within TTRs **(Fig2diii)**. This previously undetected feature shows that TTRs defined using E/L Repli-Seq contain origin activity, thus contributing to variation in TTR slope gradients. As expected, termination sites, regardless of their size, show red to blue (positive to negative) transition in OK-seq that represents the fusion of opposing replication forks in termination zones (**Fig2div,Fig2dv)**. While smaller termination sites (<100kb) may be devoid of origin activity, larger termination sites (>100kb) must contain origins to replicate so rapidly. Consistently, polarity transitions at large termination sites (>100kb) are of a lower magnitude overall than those at smaller termination sites (<100kb), suggesting the presence of increased origin activity that diminishes the polarity bias of Okazaki fragments. However, while initiation within large termination sites (>100kb) is less efficient and more distributed, the transitions from TTR to termination sites are well defined, suggesting that initiation at those transition sites (the edges of large termination sites) is more efficient. In addition, the fact that polarity transitions at termination sites (red to blue) are less uniform than those at IZs (blue to red) suggests that termination is more heterogeneous than initiation.

### TTRs are highly uniform and consistent with long uni-directional forks

Our demonstration that TTR breakages represent small IZs that disrupt TTR slope, implies that removing breakages from TTR analysis should enable more accurate estimation of fork speed within TTRs. In fact, we found that such “breakage-free” TTRs are remarkably uniform in slope and defined in replication timing within the population. In order to estimate the frequency of active origin firing within TTRs, we calculated the distribution of fork speed inferred by dividing TTR sizes by fractions of S phase traversed by TTRs in human and mouse cell lines. Assuming a 10hr S phase (the maximum for the 5 cell lines profiled), the median speed of fork progression for both human and mouse was between 1.7kb/min and 1.8kb/min (with the exception of HCT116, which has a median of 2kb/min) and agrees with the range of estimated fork speed measured directly by DNA fibre methods by others (**Fig2e**) (Pereira et al., 2017; Wilson et al., 2016). Although we cannot rule out the possibility of sequential origin firing with inter-origin distances <50kb, our limit of detection, there would need to be a nearly uniform density and firing efficiency of origins along the length of each TTR to escape detection. Neither OK-seq nor the SNS data nor single DNA fibre mapping data support such an initiation pattern in any of the cell lines analysed. We conclude that TTRs are remarkably uniform in replication speed within a population and can be accounted for by unidirectional fork movements devoid of origin firing.

### Measuring RT heterogeneity genome-wide

The strong enrichment of nascent DNA within only a few temporal windows of S phase indicates a high degree of cell to cell conservation for all kinetic features of the RT programme genome-wide. However, there are also regions where nascent DNA signal was spread over a broader temporal interval. To measure the degree of temporal variability across the genome and within the different patterns identified in **Fig1d**, we examined the relationship between two important indices of replication dynamics, Trep (the time point at which 50% of all cells have finished replicating the locus) and Twidth (the time difference between the locus being replicated in 25% of all cells and 75% of all cells) (Yang et al., 2010). Since the 16 fraction heatmap values are representative of the percentage of cells in the population that have replicated the genomic bin in each time interval, we can estimate T_rep_ and T_width_ by performing a sigmoidal fit on the column-wise cumulative sum of the bin values (**Fig3a**). Briefly, the cumulative sum for any genomic locus is calculated as such: at S phase fraction 1 (S1) the cumulative sum equals to the value the genomic bin assumes in S1, increases to the sum of S1 and S2 at the end of S2 and eventually reaches 100%, the sum of S1, S2, S3 through to S16 at the end of S phase. The value of Twidth represents the variation in replication time. For each of the features shown in **Fig 1d**, we first combined the bins within the boundaries of each feature (defined as **Fig1d**) and then measured T_width_ for the bins within each feature collectively. IZs were divided into four categories (early, early-mid, late-mid, late) as in **Fig2a**. For all cell types assayed, the T_width_ increased as S phase progressed towards mid S phase and decreased from mid S phase to late S phase (**Fig3bi,ii**). The relationship between T_width_ and T_rep_ of TTRs assumes a similar pattern to that of IZs. Variability is the highest for mid S phase for all features measured. Assuming a 10 hr S phase, by converting 16 S phase fractions into fractions of 10 hrs we found that almost all regions went from being 25% replicated to 75% replicated in 1.25-2.5 hrs. We conclude that the time at which each IZ will initiate replication is remarkably uniform within the majority of cells in the population. Remarkably, we found that all features of the RT programme, including TTRs and termination sites (<100kb), which both appear devoid of origin activity by the measurements described above, also display the pattern of increased variability during mid-S phase, characteristic of IZs (**Fig3biii,iv,v,vi)**.

**Figure 3.**
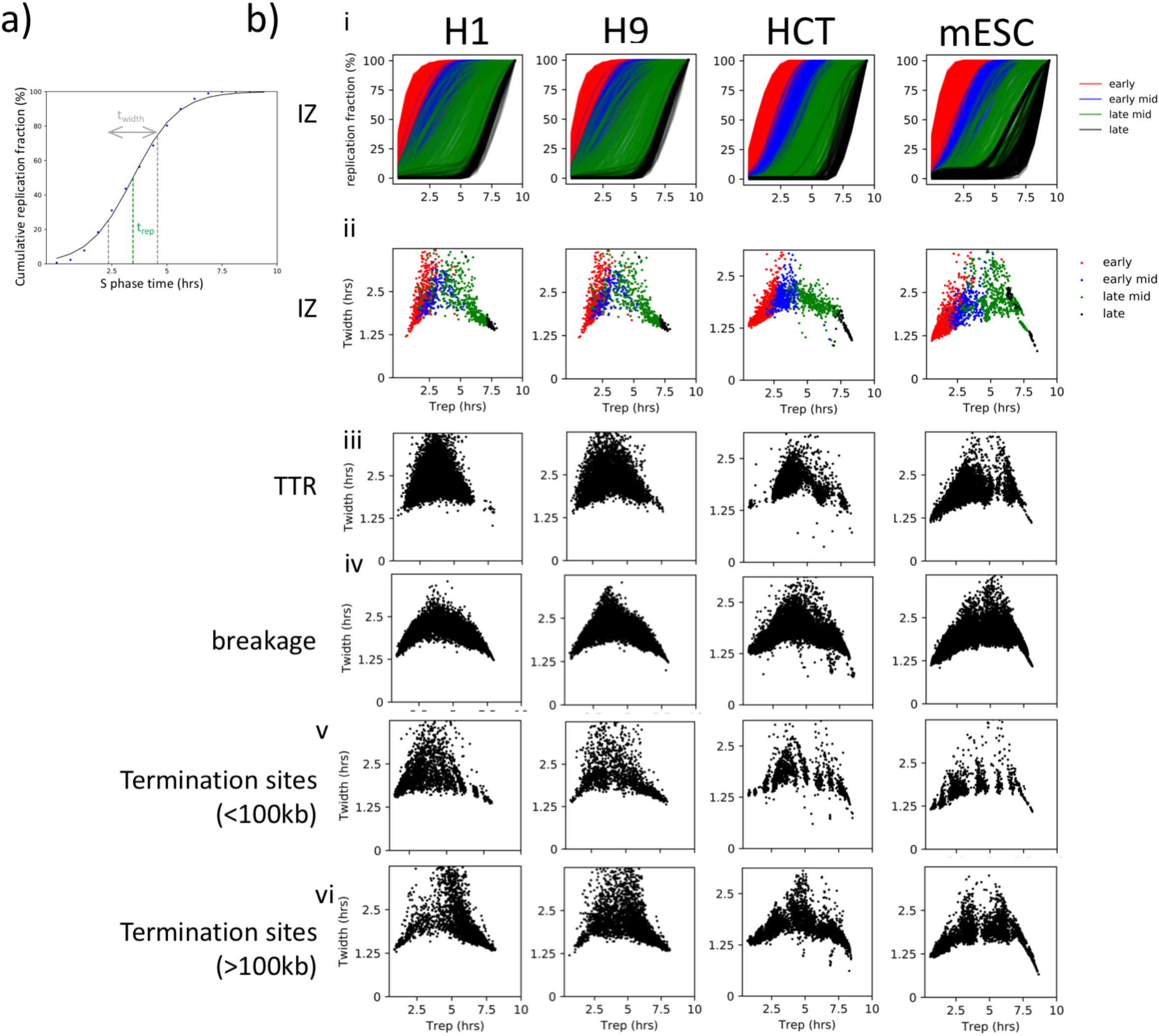
RT heterogeneity fluctuates throughout S phase and peaks in the middle of S phase. a) Schematic showing the fitting of a sigmoidal curve on the bin wise cumulative percentage replicated plotted against time into S phase. Blue points are example datapoints of cumulative percentage of replication in High-Resolution Repli-Seq heatmap. Black line represents the fitted sigmoidal curve. bi) Cumulative replication percentage for IZs of early (red), early mid (blue), late mid (green) and late (black) timing where each line represents an individual IZ. bii) Scatter plots showing the relationship between T_width_ and T_rep_ calculated from fitted sigmoidal curves for early (red), early mid (blue), late mid (green) and late (black) IZs in H1, H9, HCT116 and F121-9 mESC. biii-vi) Scatter plots showing the relationship between Twidth and Trep for TTRs (iii), breakages (iv), termination sites (<100kb) (v) and termination sites (>100kb) (vi) in H1, H9, HCT116 and F121-9 mESC.

### IZs are enriched for active histone marks and depleted of repressive marks

There have been numerous reports linking origin firing with active histone marks and active gene transcription (Fragkos et al., 2015). Centring on IZs, we performed a series of pileups showing the enrichment of histone modifications H3K4me3, H3K27ac, H3K27me3 and H3K9me3 at IZs (**Fig4a-d)**. Early, early-mid and late-mid IZs are enriched for H3K27ac, which marks active transcription start sites (TSSs) and enhancers, as well as H3K4me3, which marks TSSs. Levels of these active histone marks in the regions surrounding IZs were positively correlated with earlier replication timing. This correlation was consistent across all cell types, suggesting that these active marks are not necessary for initiation per se but correlate with early replication, possibly due to their correlation with other features such as transcription. By contrast, repressive histone marks are much more variable between cell types with respect to their enrichment at IZs or their correlation to replication timing, consistent with prior reports showing that differences in active but not repressive histone marks correlate with differences in replication timing between cell lines (Yue et al., 2014). H3K9me3 shows a slight depletion in almost all IZs except for the late ones, which are characterised by an elevated level of H3K9me3 highlighting that there is no genome-wide correlation or anticorrelation between any of the queried histone marks and IZs. In H9 hESCs, consistent with its marking of bivalent promoters and concurrence with H3K4me3, H3K27me3 is enriched at early, early-mid and late-mid IZs (**Fig4a)**. The highest concentration of H3K27me3 is found in late-mid IZs. In HCT116, however, H3K27me3 is depleted from early, early-mid and late-mid IZs with late IZs showing no depletion or enrichment **(Fig4b)**. In mouse ESCs, H3K27me3 is depleted from only early IZs; it does not show depletion or enrichment for IZs of other timing (**Fig4c)**. In mNPCs, however, H3K27me3 is mostly enriched in early IZs and depleted in late IZs **(Fig4d).** Furthermore, to define the histone associations at developmentally controlled IZs, we examined H3K27ac localisation around mESC- and mNPC-specific IZs defined in **Fig2b.** We found that H3K27ac was sharply localized within the mESC-specific IZs. This localization was also seen in mNPCs where the IZ is not active, showing that active histone marks are not sufficient for replication initiation activity. mNPC-specific IZs were enriched for H3K27ac over much broader regions and this broad enrichment was also seen in mESCs where those IZs are not active (**Supplementary Fig7).** This variable relationship between IZs and histone modifications further highlights the complexity underlying the link between replication initiation and the local chromatin environment and the absence a one to one correlation between initiation and any individual epigenetic mark.

**Figure 4.**
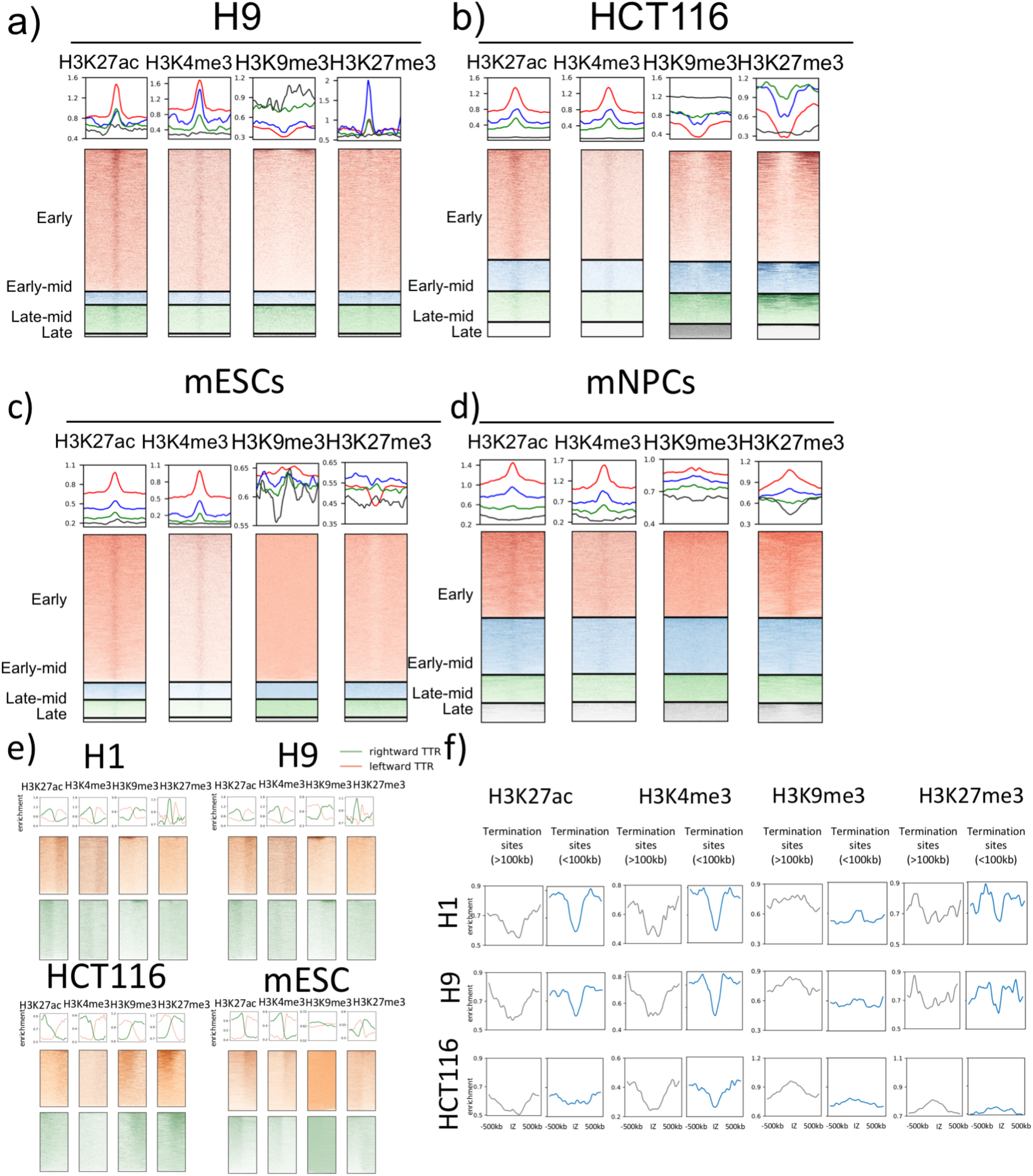
Correlation between IZs and histone modifications is cell type and timing specific. a-d) Mean line plots (top row) and heatmaps (bottom row) of H3K27ac, H3K4me3, H3K9me3 and H3K27me3 fold enrichment signal centred on early (red), early mid (blue), late mid (green) and late (black) IZs +/− 500kb in H9 (a), HCT116 (b), mESC (c) and mNPC (d). e) Mean line plots (top row) and heatmaps (bottom row) of H3K27ac, H3K4me3, H3K9me3 and H3K27me3 fold enrichment signal centred on TTRs +/− 500kb in HCT116, H1, H9 and mESC. Green and orange lines/heatmaps indicate rightward and leftward TTRs respectively. f) Mean line plots of H3K27ac, H3K4me3, H3K9me3 and H3K27me3 fold enrichment signal centred on termination sites (>100kb) (top row) and termination sites (<100kb) (bottom row) +/500kb in HCT116, H1 and H9.

TTRs, on the other hand, are depleted of active histone marks themselves and the surrounding histone modification patterns are consistent with the presence of IZs upstream and downstream of rightward and leftward TTRs, respectively (**Fig4e**). H3K4me3 and H3K27ac are enriched on the IZ side of the TTR for all cell lines. H3K9me3 and H3K27me3 shows cell type specific features that echoes their relationship with IZs. For instance, mESC IZs show no depletion of H3K9me3 consistent with its even distribution across TTRs (**Fig4c,e).** HCT116, on the other hand shows depletion of both H3K9me3 and H3K27me3 at its IZs, which is in agreement with its preferential colocalisation on the termination side of TTRs (**Fig4b,e)**. Together this shows that TTRs and IZs are differentiated by their distinct epigenetic features. TTRs lack the active marks that exist at IZs, which could potentially explain their lack of origin activity. Therefore, TTRs punctuate both chromatin marks and initiation potential.

Moreover, we show that sites of termination are associated with a paucity of active marks and enriched for repressive marks regardless of their sizes (**Fig4f).** Cell line specific differences are also observed. For instance, H3K27me3 is enriched in both types of termination sites in HCT116 yet depleted in those in H1 and H9 hESCs. Overall, despite the origin activity present in large termination sites (>100kb), they are enriched for repressive marks to the same extent as small termination sites (<100kb) that are devoid of origin activity suggesting that origins in large termination sites (>100kb) are activated independently from the enhancing effects of active marks. This supported by the observation that late IZs (Fig. 4a-d) are the most similar of all IZs to the chromatin features of termination sites. Perhaps additional mechanisms come into play late in S phase to ensure completion of replication prior to mitosis.

### Replication timing and genome architecture

Active chromatin marks and active gene transcription have been shown to correlate with high insulation on Hi-C chromatin contact heatmaps, sometimes referred to as Topologically-Associating Domain (TAD) boundaries. Consistently, we find that early and early-mid IZs, which are characterised by active chromatin marks, also correspond to loci of high level of insulation (**Fig5a**) while late IZs, which lack active histone marks, do not colocalise with increased insulation. Moreover, initiation zones identified by (Petryk et al., 2016) using OK-seq data were shown to fire preferentially at TAD boundaries and those overlap with early but not late IZs. Surprisingly, mNPCs did not conform to the correlation seen in hESCs, mESCs and HCT116 in that early IZs in mNPCs showed decreased insulation compared to IZs of later initiation timing. (**Fig4d**). In summary, the correlation of IZs to TAD boundaries is incomplete and cell-type specific, potentially driven indirectly by other factors that colocalise with IZs and share the same histone marks. It will be important to examine these correlations in other cell types.

**Figure 5.**
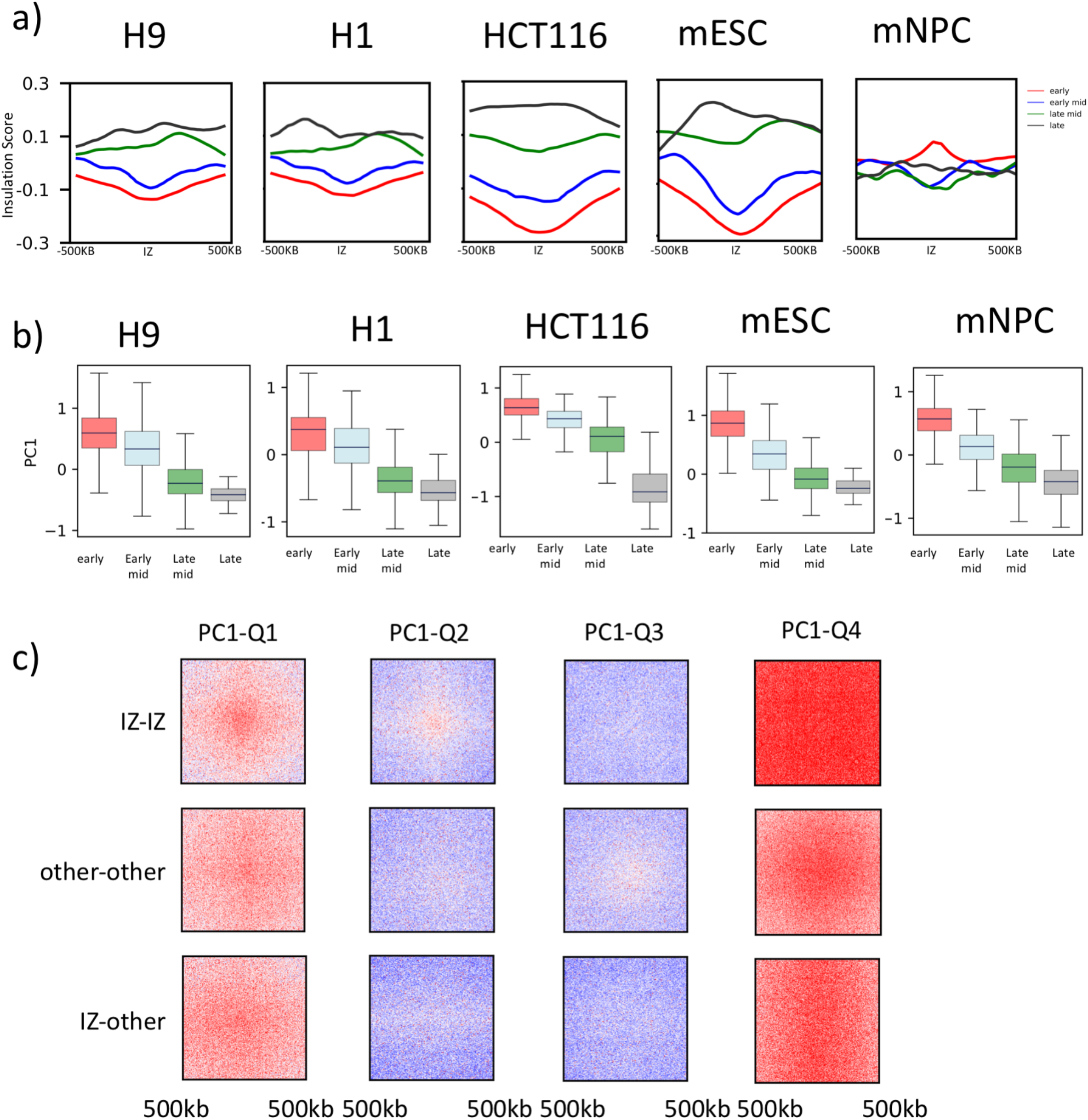
IZs of varying timing show distinct chromatin interaction preferences. a) Mean line plots of insulation scores centred at for IZs of early (red), early mid (blue), late mid (green) and late (black) timing +/− 500kb in H9, H1, HCT116, mESC and mNPC. b) Boxplot showing the distribution of PC1 values associated with IZs of early (red), early mid (blue), late mid (green) and late (black) timing in H9, H1, HCT116, mESC and mNPC. Black lines indicate medians of distribution. c) log2(observed/expected) HiC contacts for inter IZ interactions (IZ-IZ top row), non-IZ feature to non-IZ feature interactions (other-other middle row) and IZ to non-IZ feature interactions (IZ-other bottom row) with 500kb up and down-stream from the centre in HCT116. ‘Other’ is defined as all replication features in Fig1d except for IZs. Eigenvector PC1 values of HCT116 are divided into 4 quartiles. Only interactions between features of the same quartile of eigenvector PC1 value distribution are calculated for the heatmaps.

We further queried the Hi-C A/B chromatin compartment in which each of the temporal classes of IZs resides. Consistent with prior studies showing high positive correlation between RT calculated from E/L Repli-seq and Hi-C eigenvector values ((Pope et al., 2014; Ryba et al., 2010; Yaffe et al., 2010), the timing of initiation conforms to the compartment identity of the site. The earliest IZs are characterised by the most positive eigenvector values which represent the A compartment. As IZs initiate at later times in S phase, eigenvectors of these regions decrease accordingly (**Fig5b**). In contrast with histone marks, this property was consistent between all cell lines and types, underscoring the close relationship between replication initiation and chromatin compartment.

Since it has been shown that chromatin with similar histone marks shows strong intra-compartmental interactions (Rao et al., 2014), we hypothesised that IZs, which are mostly enriched for active histone marks relative to other features will also interact with each other above background in Hi-C heatmaps (**Fig5c)**. In order to ensure that any potential differences in interaction frequency are not due to the loci of interest being in different chromatin compartments, we divided the genome into quartiles according to their PC1 values calculated from Hi-C contact maps and produced pile ups of log2(observed/expected) interaction counts for IZ-to-IZ, IZ-to-other (all non-IZ features), and other-to-other for each quartile. IZs of the first two quartiles interact with one another above the background more than with other features of the same PC1 quartile **(Fig5c).** Since these early IZs are enriched for H3K4me3 and H3K27ac, IZ-to-IZ interactions could consist of functional loops such as promoter-enhancer loops and interactions between Early Replication Control Elements (ERCEs) (Sima et al., 2019). IZs in the last quartile of PC1 distribution interact significantly with each other over a larger region (>1Mb) than any other homotypic or heterotypic interaction between other replication features (**Fig5c).** Thus, these late IZs, which are contained within large termination sites and presumably contain many distributed origins, form strong intra-B compartment interactions over MB long regions. Overall, strengths of IZ-IZ interactions are consistent with their chromatin states in that the strongest interactions are formed by those IZs that overlap with functional elements marked by H3K4me3 and H3K27ac. Overall, these data show that IZs interact more frequently than with other features of similar eigenvector values, contributing to compartment formation.

### Bi-phasically replicated regions contain long transcribed genes, suggesting a link to genome fragility

Consistent with prior reports (Hansen et al., 2010), we observed biphasic patterns of replication in H1, H9 hESCs and HCT116 (**Fig6a,b**), which are defined as regions with two distinct times of replication. We first considered whether these were regions of imprinting, known to show allele-specific asynchrony (Donley and Thayer, 2013). We found that some imprinted genes show smearing in Repli-Seq heatmaps, indicative of small and stochastic differences in the RT. However, their RT differences did not reach our threshold of biphasic patterns, which required an intervening temporal interval when we could not detect replication of either allele. As a whole, the correlation between alleles at imprinted genes remained high and the majority of biphasic loci did not contain imprinted genes (**Supplementary Fig8**). Surprisingly, however, we found that the biphasic sites in human cells were associated with common fragile sites (CFS) and/or large genes (> 200kb), which are known to be correlated with genome instability and CFSs (Debatisse and Rosselli, 2019). Moreover, biphasic patterns were cell-type specific, as are CFSs (Debatisse and Rosselli, 2019; Letessier et al., 2011). We identified 340 biphasic replication sites in H1/H9 hESCs, 144 of which (42%) were CFSs (annotated in other cell lines) and 116 that have not been shown to be fragile in any cell line but contained large genes (> 200kb). HCT116 contained 26 biphasic sites. CFSs have been mapped cytogenetically in HCT116 (Le Tallec et al., 2013) and 9 out of 10 of those that map to autosomes displayed biphasic replication patterns, validating the fragility of sites detected by High-Resolution Repli-Seq (**Fig6 a,b,c**). Interestingly, the single undetected CFS in HCT116 was fragile in only 1.3% of metaphases, while the other 9 were present in more than 1.6% of metaphases, suggesting an approximate sensitivity of High-Resolution Repli-Seq to detect CFSs. 10 of the remaining 17 HCT1216 biphasic sites that did not overlap with mapped CFSs nonetheless contained large genes or overlapped with CFSs mapped in other cell types and could be potential novel fragile sites. We found that biphasic regions are marked by active histone marks such as H3K4me3 and H3K27ac, which suggests that they are potentiated for transcription or under active transcription (**Fig 6d**). We compared the levels of transcription of large genes overlapping with biphasic loci against those of large genes genome-wide in HCT116, H9 and H1 hESCs and found the medians of Transcripts per million (TPM) distribution were higher in biphasic genes, suggesting that they are generally expressed (**Fig6e**). In conclusion, biphasic patterns provide a replication signature of transcribed long genes and CFSs.

**Figure 6.**
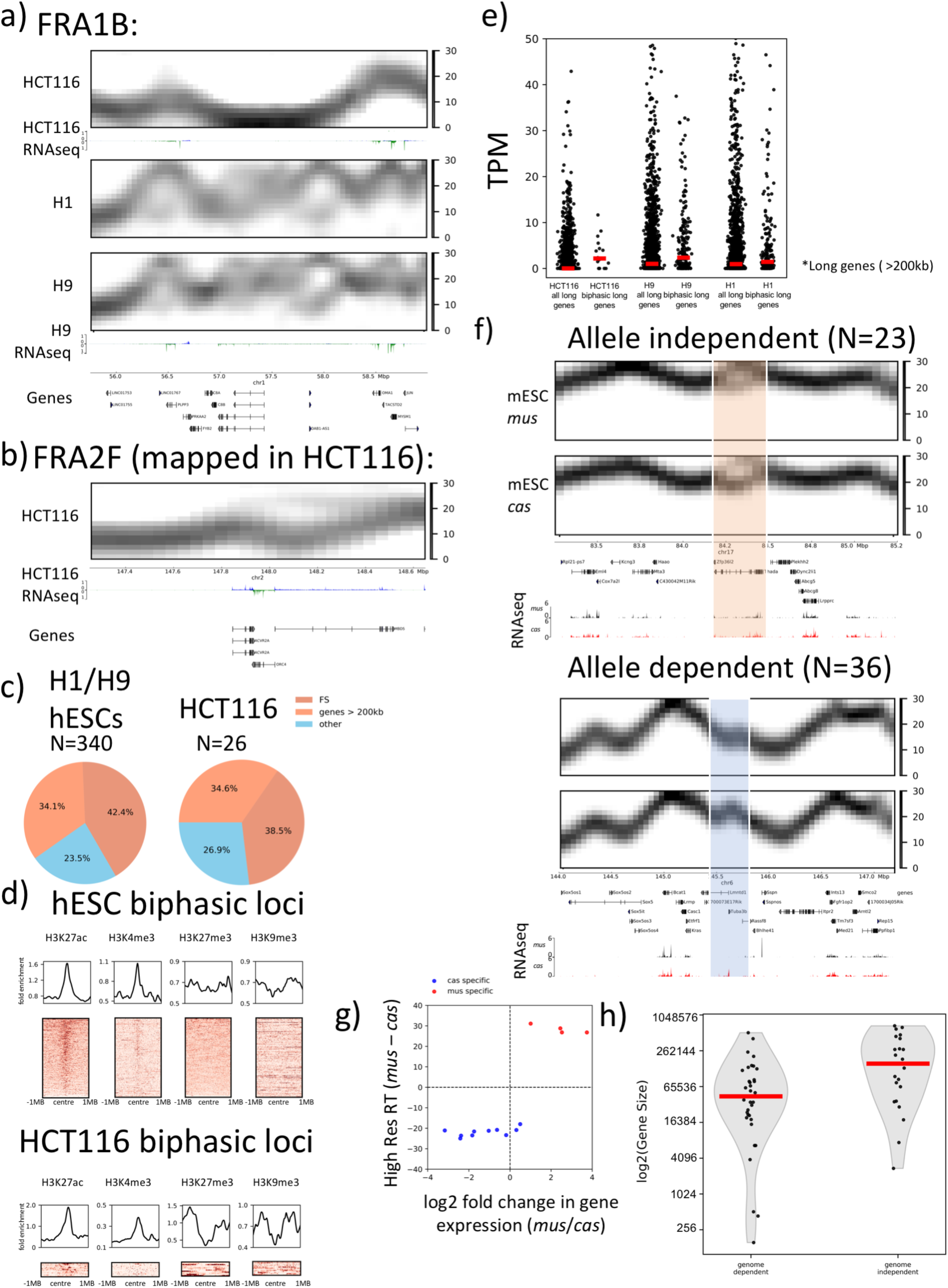
Bi-phasically replicated loci overlap with long genes that suggest genome fragility. a) High resolution Repli-Seq heatmaps and RNA-seq profiles for FRA1B (chr1:55,750,000 – 59,000,000), in HCT116, H1 and H9 hESCs. b) High resolution Repli-Seq heatmaps and RNA-seq profiles for FRA2F (chr2:147250000-148650000), a mapped HCT116 CFS, which is replicated biphasically in HCT116. c) Fractions of biphasic loci overlapping with annotated CFS and long genes (>200kb) in hESCs and HCT116. d) H3K27ac, H3K4me3, H3K9me3 and H3K27me3 ChIP-seq fold enrichment signal centred on biphasic loci in H9 and HCT116 cell lines e) Distribution of TPM of genes >200kb genome wide or genes >200kb overlapping with biphasic regions in HCT116, H9 and H1 hESCs. Red lines represent the medians of TPM. f) F121-9 mESC allele resolved Repli-seq heatmaps and RNA-seq tracks at genome independent and dependent loci. Orange highlight indicates the genome independent biphasic site in chr17, which actively transcribed on both *mus* and *cas* alleles. Blue highlight indicates the genome dependent biphasic site in chr6, which is replicated earlier in *cas* allele accompanied by active transcription. g) Correlation between log2 (gene expression differences) and RT differences of *mus* over *cas* at genome dependent biphasic sites. h) Distribution of log2(gene size) of genes overlapping with genome dependent or genome independent biphasic sites. Red lines represent the medians of log2(gene size).

To determine whether bimodality arose from replication asynchrony between maternal and paternal alleles or replication asynchrony on a population level, we plotted separate maternal and paternal Repli-Seq heatmaps in hESC line H1 in which SNPs have been phased. The high similarity between heatmaps for the locus shown in **Fig6a** did not support an allelic difference but the sparsity of SNPs in human cells posed a challenge for a statistically sound conclusion. For this reason, we examined the F121-9 cell line, which allows effective allele parsing due to its high SNP density. There are 59 sites in F121-9 mESCs that were found to be biphasically replicated, 36 of these were found to be due to allelic differences. The remaining 23 sites were found to be allele independent; the biphasic patterns were retained in heatmaps for individual alleles (**Fig6f**). Moreover, allele-dependent bimodality was associated with differential gene expressions between alleles. In the example shown in **Fig6f**, the earlier replicating locus on the *cas* allele coincides with higher transcription of the Tuba3b gene on the same allele. The same trend holds true genome wide in that the earlier replicating locus on either allele was associated with higher levels of gene transcription within the locus (**Fig6g**). In contrast to allele-independent loci, allele-dependent loci showed no preferential affiliation with large genes. Indeed, we find that larger genes preferentially overlap with allele-independent biphasic sites than allele dependent (**Fig6h**). We conclude that biphasic replication at allele-dependent and independent loci are orchestrated by distinct mechanisms, with allele-independent (random) asynchrony associated with large expressed genes that in humans are linked to CFSs and allele-dependent asynchrony potentially encoded by polymorphic differences between the genomes.

## DISCUSSION

The apparent paradox between highly stochastic replication origin usage in mammalian cells and the considerably more deterministic nature of their RT patterns has been the focus of many studies (Rhind et al., 2010). By using a short nascent DNA labelling time and increased number of S phase fractions we have produced Repli-Seq profiles with exquisitely high temporal resolution, revealing finely choreographed temporal structure in genome replication that is remarkably uniform between cells in a population. The data reveal discrete initiation zones at 50kb resolution that initiate at various time throughout S phase. Bi-directional replication forks can be detected emanating from all temporally distinct IZs but those with different initiation times are differentiated by characteristics such as SNS origins, chromatin mark enrichment or Hi-C interaction strength and breadth. We show that when sparse regions of initiation are filtered out of TTRs, the remaining “true TTRs” can be accounted for by single, very long, unidirectional replication forks. When all features of the genome consistent with initiation are compiled, we estimate that 72% of 50kb segments of the genome harbor initiation sites. Finally, our data reveal previously undetected regions of replication bimodality some of which are linked to allelic polymorphisms and some that are sites of random asynchrony, the latter of which overlap with CFSs or large genes (>200kb) associated with genome instability.

We identified IZs as peaks in high resolution Repli-Seq heatmaps throughout S phase, which are subsequently cross referenced with origin mapping methods such as SNS-seq and OK-seq. Early IZs show enrichment in SNS-seq while late IZs do not. On the contrary, IZs of all timing show the corresponding directionality switch in fork polarity in OK-seq data suggesting that high resolution Repli-Seq and OK-seq identify initiation events of similar characteristics. Both methods can detect broad zones of initiation containing origin clusters as well as zones with one or a few highly efficient sites that are detected at higher resolution by the SNS-seq method and so they are less sensitive to the decrease in individual origin efficiency as S phase progresses. The enrichment of SNS-seq signal in early IZs is also consistent with the preferential colocalization between early IZs with active histone marks, a defining characteristic of origins identified through SNS-seq. In accordance with the depletion of SNS-seq signal in late IZs, late IZs are also devoid of active histone marks. We also show cell-type specific correlation between histone modifications and origin firing thus, while a permissive chromatin environment might enhance the probability of firing and the formation of origin clusters manifesting as IZs, origins can fire in an environment defined by repressive histone marks.

In addition to IZs, we defined TTRs in high resolution Repli-Seq heatmaps and approximated the fork speed that would give rise to the slopes seen in TTRs in a 10hr long S phase. The approximated speeds are in good concordance with the range of fork speeds obtained through DNA combing and other methods. HCT116 TTRs has a slightly higher median speed (2kb/min) than stem cells, which agrees with a prior study reporting higher TTR speed in cancer and differentiated cells than in stem cells. The primary mode of replication in TTR has been controversial, with some reports consistent with passive uni-directional fork movement (Hiratani et al., 2008) and others claiming that slopes of TTRs are too fast and must contain sequentially activated origins (Guilbaud et al., 2011). Our results demonstrate that prior datasets were not able to resolve true TTRs from small inefficient zones or shoulders of origin activity (breakages). High resolution Repli-Seq allowed us to remove these breakages in the slopes, thus filtering for only true TTRs with continuous slopes. Overall, the human and mouse stem cell lines show very similar median TTR speed that is in the range of previously measured singular fork velocity in these cell types. Importantly, with breakages removed, slopes are highly uniform within a cell type.

High resolution Repli-Seq heatmaps reveal biphasic replication patterns that are previously undetected in E/L repli-seq. Biphasic sites can be either allele dependent or allele independent. Allele-dependent bimodality is typically associated with a higher gene transcription rate in the earlier initiating allele. However, not all transcriptional differences result in replication asynchrony and vice versa hence ruling out an absolute correlation between active transcription and replication initiation. Allele-independent bimodality is enriched for large genes and CFSs. Previous studies have concluded that CFSs exhibit delayed replication by using E/L Repli-Seq (Pelliccia et al., 2008). However, E/L Repli-Seq produces an averaged RT profile and a read enrichment in both early and late S phase fractions would render a locus seemingly mid-replicating. Through High-Resolution Repli-Seq, we have found a specific biphasic replication signature associated with CFSs and may provide a means to predict novel CFSs. The precise cause for biphasic replication at large genes is elusive. We hypothesise that the biphasic patterns are due to heterogeneous random mono-allelic levels of transcription at these large genes in the cell population. In the fraction of cells where the gene is actively transcribed or poised for transcription, the open chromatin induced by transcription may be conducive for activation of nearby origins, which facilitates earlier replication of the locus. In the fraction of cells where the gene is not actively transcribed, it is passively replicated by fork passage from neighbouring IZs and is sufficiently delayed that it can be completely resolved, creating a biphasic pattern in the High-Resolution Repli-Seq heatmap.

## ACKNOWLEDGEMENTS

We thank D.Janssens and S.Henikoff for sharing H1 Cut-and-Run datasets with us prior to publication, M.K. Parsi, J.Gibcus, S.Venev, B.A.Oksuz, R.Maehr and J.Dekker for sharing H1 Hi-C datasets with us prior to publication. We are grateful to N.Rhind, J.Ma, B.V.Steensel for their helpful comments on the manuscript.

## MATERIALS AND METHODS

### Cell Culture

WA01 and WA09 cells were cultured in mTeSR1 (StemCell Technologies #85850) on hESC-qualified Matrigel (Corning #354277)-coated dishes according to WiCell instruction. For the maintenance and expansion, cells were detached using ReLeSR (StemCell Technologies #95872). When the cell reached approximately 70% confluent, cells were pulse-labeled with 400uM BrdU for 30 minutes. To obtain single-cell suspension easily, the BrdU-labelled WA01 and WA09 cells were harvested using Gentle Cell Dissociating Reagent (StemCell Technologies #07174). HCT116 cells were cultured in McCoy’s 5A medium supplemented with 10% FBS. When the cells were approximately 70% confluent, cells were pulse-labelled with 400uM BrdU for 30 minutes. The BrdU-labelled HCT116 cells were harvested using trypsin-EDTA. F121-9 mouse embryonic stem cell was cultured in 2i media on gelatin-coated dish as described in (PMID:29735606). F121-9 differentiation to NPC was performed using RHB-A (TakaraBio #Y40001) as described in (PMID:29735606) for 10 days. F121-9 ESC and NPC were pulse-labelled with 400uM BrdU for 30 minutes and harvested using ESGRO Complete Accutase (Millipore Sigma #SF006). NPC differentiation was confirmed by 1) cell morphology, 2) qPCR for marker genes; Oct4 (ESC marker), Sox1 (NPC marker), Ptn (NPC marker), Nestin (NPC marker).

### E/L Repli-seq library preparation and sequence processing

The experiments and analyses of E/L Repli-Seq were carried out as described in (Marchal et al., 2018). Briefly, the libraries were sequenced on Hi-Seq 2500. The fastq reads were mapped to human genome hg38 or mouse genome mm10 with the parameters --no-mixed, --no-discordant. PCR duplicates were removed using samtools rmdup. Log2 E/L ratio was calculated for 50kb bins. The final profiles were Loess smoothed and quantile normalised using all profiles used in this paper.

### 16 fraction high resolution Repli-seq processing and library preparation

The BrdU labelled cells were fixed in 70% ethanol and stained with propidium iodide as described in (Marchal et al., 2018) then sorted by BD FACSAria SORP according to the DNA content. Eighty thousand cells were collected for each fraction. In order to obtain reproducible sorting windows easily, the region from G1 peak to mid-point between G2 peak and the end of G2 was equally sliced into 16 to make S1-S16 fractions (WA01, WA09). As we found S15-S16 fractions did not contain detectable BrdU-labelled DNA, in the later experiments, the region from G1 peak to G2 peak was equally sliced into 16 to make S1-S16 fractions (HCT116, F121-9 ESC and NPC). Any cells on the left of G1 peak were considered non-replicating and collected as G1 fraction. From cells in each fraction, total genomic DNA was extracted. DNA from S1-S16 fractions were made into libraries as described in (Marchal et al., 2018) with the following modifications: after the genomic DNA was sheared and adaptors were ligated, the entire adaptor-ligated DNA from 80K cells was used for one BrdU-immunoprecipitation (using 0.5ug of anti-BrdU BD #555627 and 20ug of anti-mouse IgG Sigma #M7023) since the smaller amount of BrdU-labelled DNA was anticipated due to the short pulse label (WA01, WA09) then proceeded to purification and indexing. For HCT116, F121-9 ESC and NPC, adaptor-ligated DNA from 80K cells was first incubated with 0.5ug of anti-BrdU in 100uL PSBT (0.137M NaCl, 0.0027M KCl, 0.01M Na2HPO4, 0.0018M KH2PO4, 0.1% Tween 20) for 20 minutes at room temperature, then the BrdU-DNA/anti-BrdU complex was captured by 2uL of Dynabeads Protein G (Thermofisher #10003D) directly added this reaction for 20 minutes at room temperature. The BrdU-DNA/anti-BrdU/Protein G bead complexes were washed with 200uL of PBST for 3 times (5 minutes each) before the release of BrdU-DNA by Proteinase K digestion and purification described in (Marchal et al., 2018) before library indexing.

### Sequencing, Mapping and normalisation of High-Resolution Repli-Seq Data

Repli-Seq libraries were sequenced on Hi-Seq 2500. Reads were aligned to human genome hg38 or mouse mm 10 using bowtie-2 with the same parameters as those used for E/L Repli-Seq. Reads per million (RPM) was calculated with 50kb bin size for BrdU pull-down libraries of each S phase fraction as well as G1 control. The log2 ratio between RPM of BrdU pull-down and that of G1 WGS was calculated for each S phase fraction, which was subsequently used to construct a matrix consisting of 16 rows where each row represented an S phase fraction ranging from S1 to S16 and each column represented a 50kb genomic bin. 50kb genomic bin was chosen due to the following considerations: assuming a fork speed of 1.8kb/min, 30mins of BrdU labelling would have enabled incorporation of the analogue in at least 50kb DNA per fork. Therefore, we estimate the technical limit of resolution to be approx. 50kb. Bins with values below zero i.e. bins that were smaller than the corresponding ones in G1 WGS were converted to zero. Sex chromosomes were removed and excluded from further analyses. The Repli-Seq heatmap matrix was smoothed by applying a gaussian filter with sigma of 1:

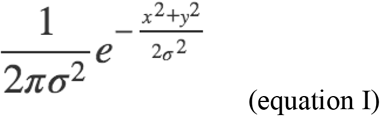

Sigma of 1 was chosen because it was the minimum kernel size so that oversmoothing could be avoided. Effectively, each genomic bin is smoothed using the values of its 8 neighbouring bins: 2 bins upstream and downstream of the target bin in the same S phase fraction, 3 bins in the previous S phase fraction and 3 bins in the ensuing S phase fraction. For the bins in the first and the last S phase fractions, the Repli-Seq heatmap maxis is padded column-wise with the first and last S phase fractions respectively. Subsequently, assuming that all bins should finish replicating at the end of S phase and should therefore be given equal weight when summing column-wise across all 16 fractions each column was assigned a total arbitrary value of 100 and Gaussian smoothed value for the bin was substituted by its original value divided by the column sum multiplied by 100.

### HiC analyses

HiC datasets are downloaded from GEO as stated in (table I) and aligned to hg38 or mm10 using HICUP (https://www.bioinformatics.babraham.ac.uk/projects/hicup/). The raw bam files are converted to.cool files using cooler and raw contact matrices are normalised either using ICE available in cooler or distance normalised as log2 (observed/expected) contacts where the expected contacts are calculated as diagonal sum divided by the number of valid bins in the diagonal. Eigenvector decomposition is performed using the cooltool package (https://github.com/mirnylab/cooltools) and ranked using GC content. Insulation scores are calculated using the cooltool package on.cool files at 5k bin size and moving diamond window size of 500kb.

### ChIP-seq analyses

ChIP-seq and Cut-and-Run reads were downloaded from GEO as stated in Table I and aligned to hg38 or mm10 using bowtie2. The bam files from alignment were used as input for MACS2 to call peaks and generate fold enrichment bigwig files, which were used subsequently for heatmap generation in the context of IZ alignment. These alignment heatmaps were generated by constructing a matrix with each row representing an IZ centred on IZ centres and each column representing a genomic bin. The values in the matrix represent the ChIP-seq fold enrichment signal at the defined genomic distance from the centre of the region of interest.

### SNS-seq and Okazaki fragment seq

The sources of SNS-seq raw reads are stated in Table I. SNS-seq was aligned to hg38 using bowtie-2 and further processed according to (Smith et al., 2016). The source of OKseq signal bigwig files and called OKseq initiation zones for mESCs is stated in Table I. The files were converted to mm10 using UCSC utility liftOver.

### Defining features in high resolution Repli-Seq: IZs, TTRs, breakages, small termination sites (<100kb) and large termination sites (>100kb)

Features were identified through the clustering algorithm BIRCH implemented in the python package scikit-learn (https://scikit-learn.org/stable/index.html). Briefly, BIRCH algorithm was applied to the normalised Repli-Seq heatmap so that each column was assigned a predicted label depending on the cluster the column belonged to. Features were identified depending on the S phase fractions where the maxima of the cluster centres were located relative to the neighbouring bins. IZs were identified as consecutive bins whose maxima of the predicted cluster centres were within the same S phase fractions that were also earlier than their neighbouring bins both upstream and downstream outside of IZs. Leftward TTRs were identified as consecutive bins whose maxima were within S phase fractions that were earlier than their upstream neighbour and later than their downstream neighbour. Rightward TTRs were identified as consecutive bins whose maxima were within S phase fractions that were earlier than their downstream neighbour and later than upstream neighbour. Breakages in TTR were identified as consecutive bins contained within TTRs whose maxima of the predicted cluster centres were within the same S phase fractions. Small termination sites (<100kb) were identified as 1 or 2 50kb bins whose maxima of the predicted cluster centres were within the same S phase fractions and were later than their upstream and downstream neighbouring bins. Large termination sites (>100kb) were identified as consecutive bins (>2) whose maxima of the predicted cluster centres were within the same S phase fractions and were later than their upstream and downstream neighbouring bins.

### Timing heterogeneity estimation

The degree of heterogeneity of replication timing of IZs and TTRs was estimated by performing a sigmoidal fitting on the column wise cumulative replication percentage. Briefly, the sigmoidal function

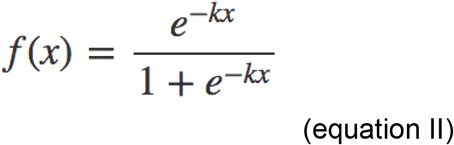

was fitted using curve_fit function in scipy, which minimises the mean squared error. T_rep_ used in the timing-variation measurement is f(x) when x is 0.5 which means the bin is 50% replicated in the cell population and T_width_ used in the timing-variation measurement is f(0.75) – f(0.25) which is the time difference between 75% replicated and 25% replicated for any genomic bin.

### Identification of biphsically replicating regions

Biphasically replicating regions are characterised by the presence of two maxima in the column-wise cumulative sum for any bin in normalised Repli-Seq heatmaps. Therefore, continuous bins that had two maxima in their column-wise cumulative sum were merged and identified as biphasic regions.

### *RNA-seq* analyses

RNA-seq data for HCT116 and H9 were downloaded from Encode (**Table I**) and aligned to hg38 using STAR to generate bigwig signal files and reads per gene counts. mESC RNA-seq was downloaded from 4DN data portal (**Table I**) and aligned to mm10 using STAR. Allele parsing was performed on parsed bam files using SNPsplit with the SNP VCF file downloaded from https://www.sanger.ac.uk/science/data/mouse-genomes-project. TPM is generated based on the output SAM files from STAR using RSEM (https://deweylab.github.io/RSEM/).

### CFS database

Genomic coordinates of human CFSs used to overlap with bi-phasically replicating loci were from https://webs.iiitd.edu.in/raghava/humcfs/

### Code availability

All bash and python codes used in this manuscript are available upon request and Github links can be found on http://replicationdomain.com.

## Supplementary Figure legends

**Figure1 Validation of BrdU pull-down using alpha- and beta globin by qPCR.**

**Figure2 Schematics showing the normalisation of Repli-Seq heatmaps.**

**Figure3 RPM of each S phase fraction is corrected using G1 WGS.** a) Cumulative RPM distribution of each S phase fraction compared with that of G1. b) cumulative fold enrichment of each S phase fraction after correcting using G1 WGS.

**Figure4 Correlation heatmaps showing concordance between High-Resolution Repli-Seq datasets of human and mouse cell lines.**

**Figure5 OK-seq IZs are primarily early replicating.** a) High-Resolution Repli-Seq pile-up heatmap for called mESC OK-seq IZs. b) Percentage of replication of called mESC OK-seq IZs in each High-Resolution Repli-Seq S phase fraction.

**Figure6 SNS-seq signal centred around rightward (darkgreen) and rightward (orange) TTRs in HCT116, H9 and mESCs.**

**Figure7 mNPC (salmon pink) and mESCs (sky blue) H3K27ac ChIP-Seq signal +/− 1MB around IZs that are mESC specific (left panel) or mNPC specific (right panel).**

**Figure8 Scatter plots showing the correlation between normalised read den-sies on *mus* and *cas* alleles in Repli-Seq heatmaps at imprinted loci for fractions S1-S16 in mESC (left) and mNPC (right).**

