## Supplementary Figures for "High Resolution Repli-Seq defines the temporal choreography of initiation, elongation and termination of replication in mammalian cells"

### Supplementary Fig 1 Zhao et al.,

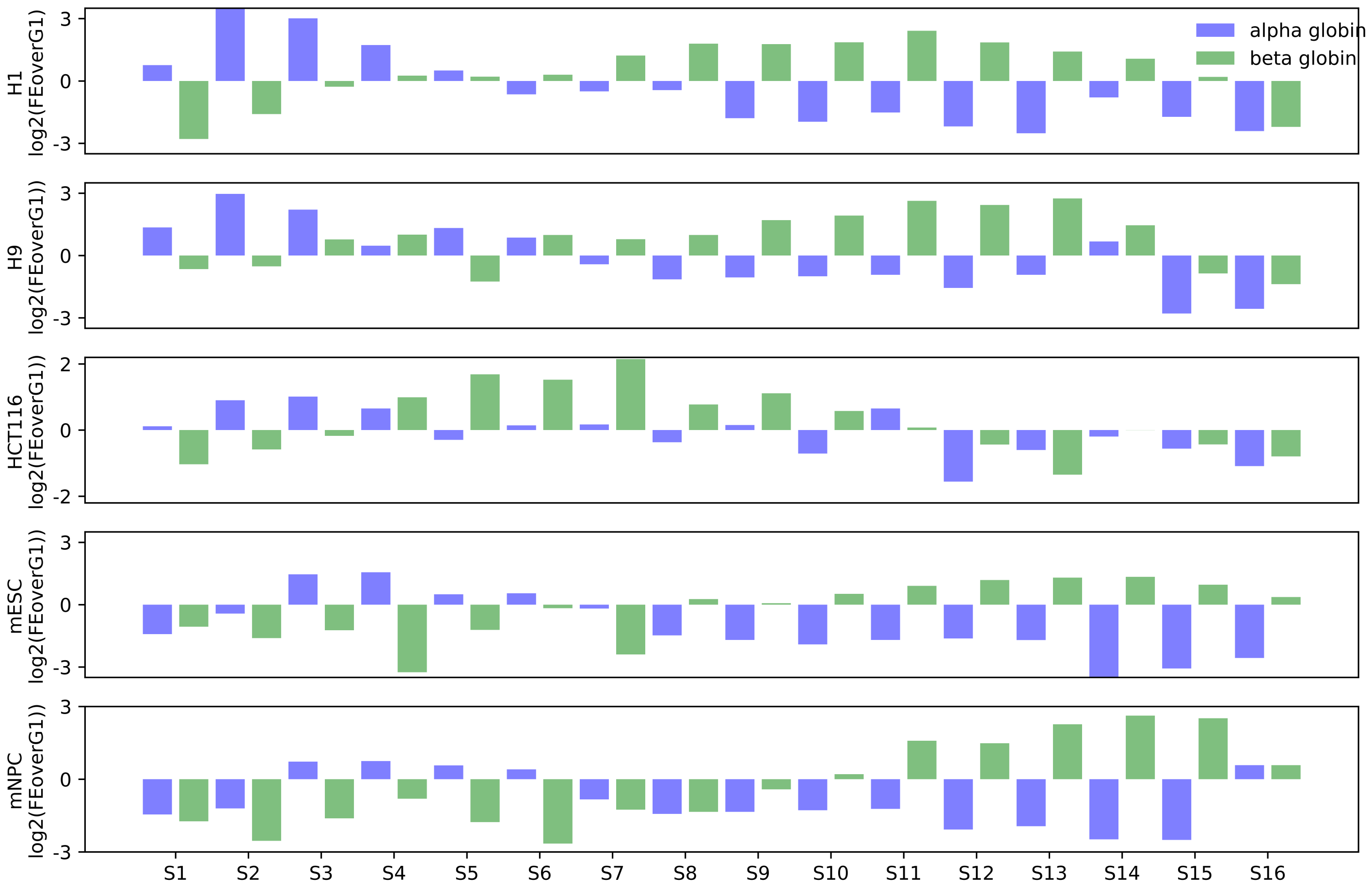

**Supplementary Fig2 Zhao et al.,**

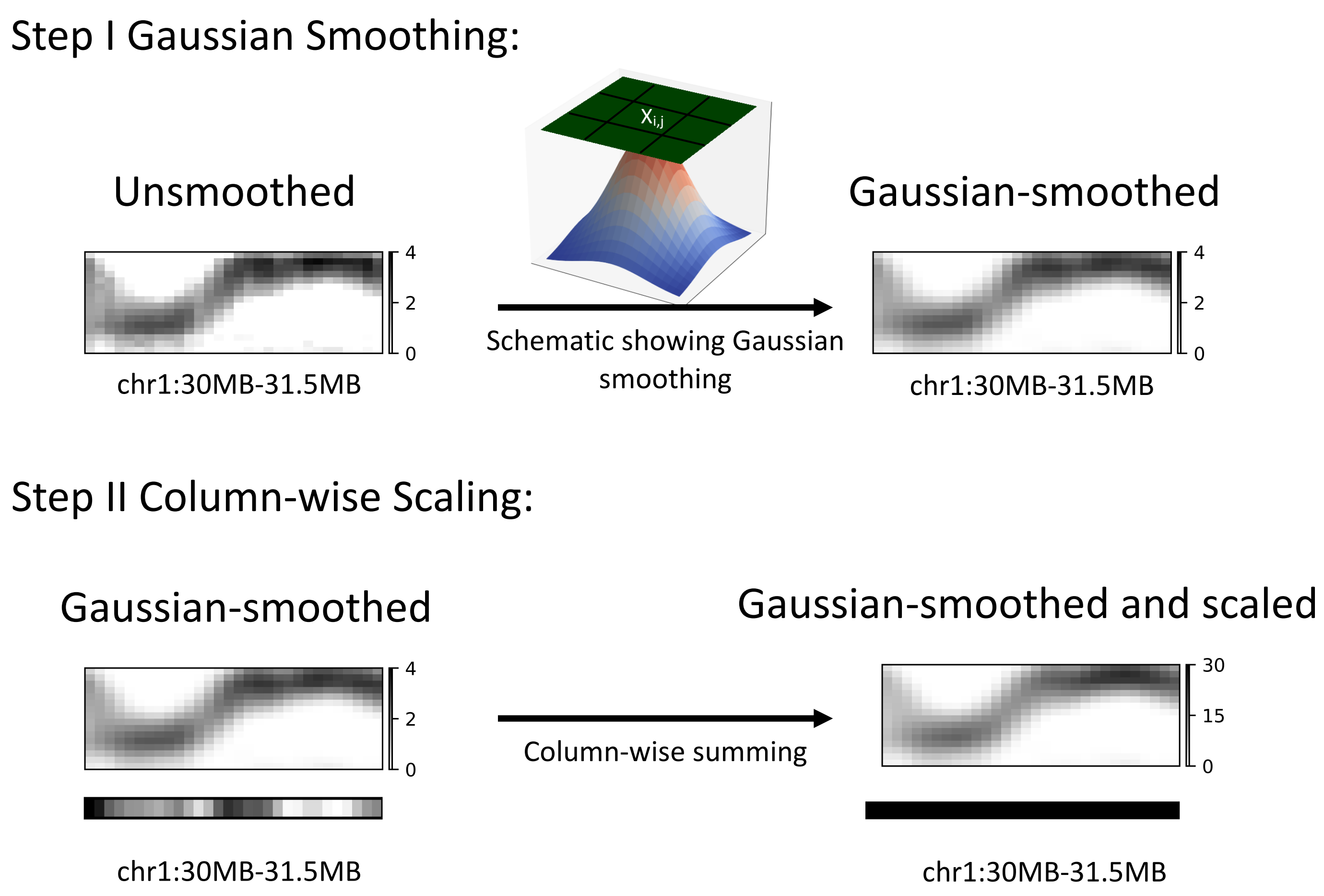

### Supplementary Fig3 Zhao et al.,

a)

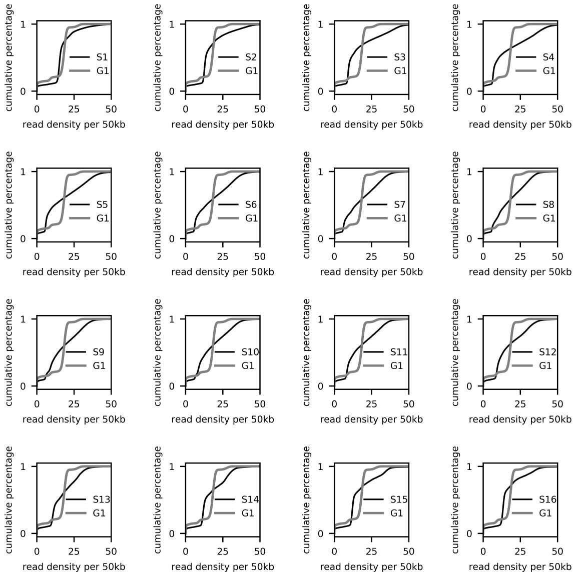


b)

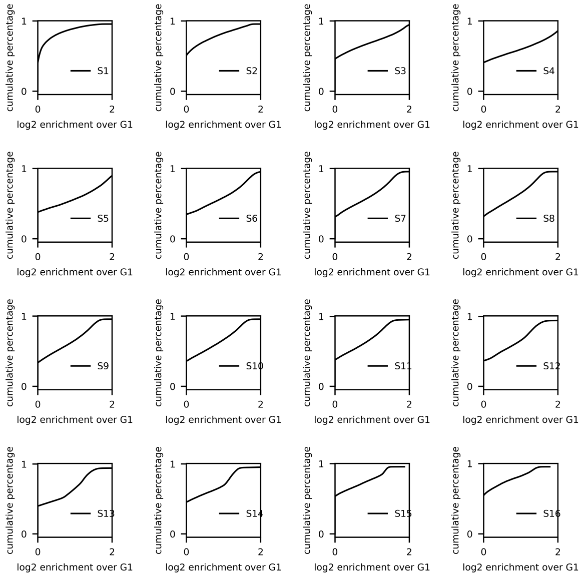

**Supplementary Fig4 Zhao et al.,**

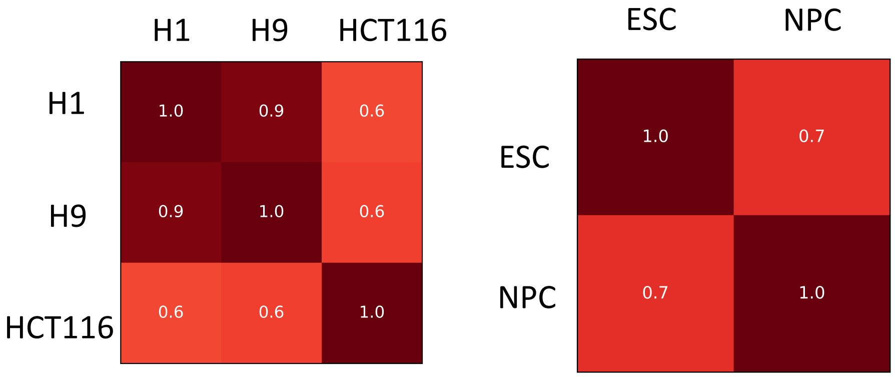

### Supplementary Fig5 Zhao et al.,

a)

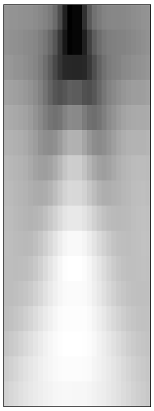

b)

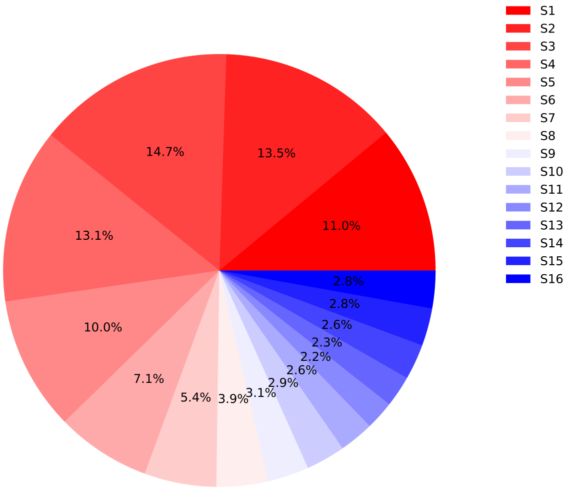

### Supplementary Fig6 Zhao et al.,

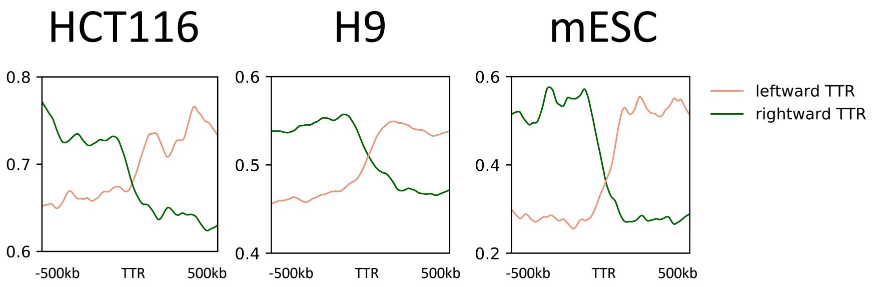

### Supplementary Fig7 Zhao et al.,

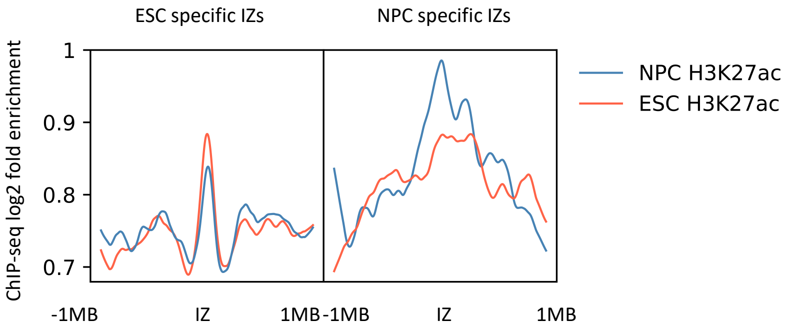

### Supplementary Fig8 Zhao et al.,

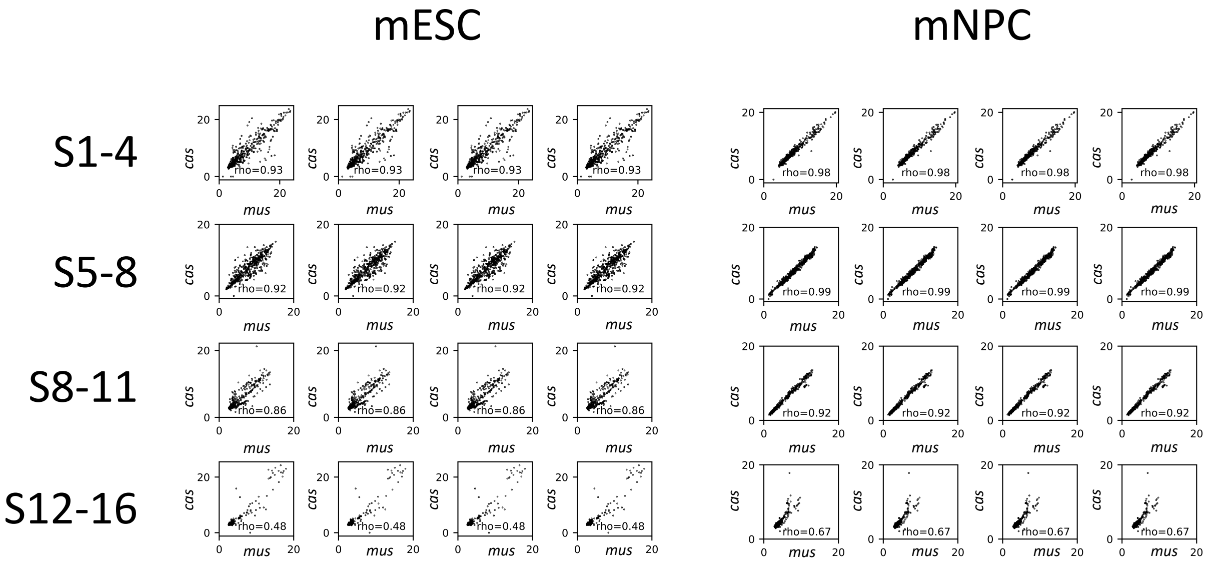
